# Quasi-static force requirements are not sufficient to explain arolium engagement in climbing Argentine ants

**DOI:** 10.64898/2026.06.29.735413

**Authors:** Yakun Cao, Andrew Chacon, Agasthya Valluri, Liam O’Connor Mueller, Nick Gravish

## Abstract

Argentine ants (*Linepithema humile*) utilize adhesive pads (arolia) to climb smooth surfaces. Previous research found that ants can adjust their individual arolium engagement according to their locomotion mode. However, it remains unclear how they distribute arolium engagement across multiple limbs to climb effectively, and how arolium engagement varies within a climbing step. As the arolium is a well-known adhesive organ, we hypothesized that engagement across different legs is distributed according to the normal forces required for balancing the body during climbing. To test this, we measured Argentine ants’ arolium engagement on a vertical glass surface using a Frustrated Total Internal Reflection (FTIR) sensor and compared it to the required normal forces from a quasi-static model. Contrary to the required normal force, the measured arolium engagement was asymmetric between upward and downward climbing, and changed over time. Our results indicated that the quasi-static force requirements are not sufficient to explain arolium engagement in climbing Argentine ants, and suggested that other factors, such as body dynamics, ants’ anatomy and behavioral preferences, should be included.

## Introduction

Some ant species, such as Argentine ants (*Linepithema humile*), can navigate smooth artificial surfaces such as glass windows and metal tables to forage [1–3]. The key anatomical feature enables this ability is the arolium [4,5], a specialized adhesive pad located on the 5th tarsomere of ants’ legs between the two claws [6,7]. To understand how ants can navigate on smooth substrates, it is important to explore how ants use their arolia.

Arolium can help ants to generate adhesion forces more than 40 times of their body weight [8]. However, ants do not always engage their arolia [9] and often avoid the full engagement [10], likely because the increased engagement can hinder rapid disengagement. This is supported by the previous findings that when exposed to strong detachment forces such as carrying weights while walking upside down [10], resisting puffs of air, or holding on to a rotating drum [8], ants walked very slowly or stayed motionless. Given the impact of full arolium engagement on their mobility, to perform movements efficiently, ants may carefully adjust the adhesive contact of their arolia to achieve a balance between generating sufficient adhesion force and maintaining locomotor speed.

Previous studies provided evidence that ants can adjust the arolium engagement [6] and disengagement [11], as well as the adhesive contact area [10] according to their locomotion mode. When Weaver ants walked upright, their 5th tarsomeres were mostly lifted with the arolia disengaged, while their arolia would adhere to the top surfaces when walking upside down [9]. Moreover, when a load was added to the upside-down ants, their arolia’s contact areas increased [10].

The prior findings were obtained at the level of single arolium [6,9,10]. However, climbing often involves multiple feet in contact with the surface at the same time. Therefore, to understand how Argentine ants achieve efficient climbing on smooth substrates, it is important to determine how they use their arolia across different legs. Previous work has shown that arolium engagement is typically present when a leg generated adhesion force [9]. Thus, we hypothesize that Argentine ants distribute their arolium engagement on different legs according to the force requirement in the normal direction. Specifically, we expect Argentine ants to engage their arolia on legs required to generate adhesion forces, and to disengage the arolia on legs required to generate compression forces. Furthermore, the leg with the higher required force should exhibit more intense arolium engagement.

To test our hypothesis, we first need to determine the required normal forces on stance-phase limbs. Previous research demonstrated that the normal force pattern of geckos—adhesion forces generated by legs above Center of Mass—could be explained by a quasi-static sagittal-plane force model [12,13]. Additionally, similar normal force patterns have been observed in weaver ants [9]. These findings suggested that a quasi-static force model can be a simple but reasonable choice. Additionally, if the arolium usage is impacted by the body dynamics, the quasi-static model could serve as a null model to compare with. As ants often climb with three feet touching the ground [14,15], we proposed a 3D tripod quasi-static force model to predict the normal forces.

We then tested our main hypothesis—whether Argentine ants distribute arolium engagement according to normal force requirements—by checking whether the measured engagement on a glass surface during vertical climbing matched the normal force pattern predicted by the quasi-static model. Using a Frustrated Total Internal Reflection (FTIR) sensor, we detected which arolia on the stance-phase legs were engaged with the glass according to the occurrence of illumination and measured the light flux from each illuminated contact as an indicator of engagement intensity. We performed three specific analyses on Argentine ants upward and downward climbing: (1) we checked whether the arolia required to generate adhesion forces were engaged; (2) we examined whether the ratio of arolium engagement intensity between different legs was positively associated with the ratio of their model-predicted required adhesion forces; (3) given the time-invariance of quasi-static force requirements, we tested if the engagement status of each arolium stayed the same during each step.

## Materials and Methods

### Study animals

Argentine ants (*Linepithema humile*) with surrounding soil were collected from 8 locations across University of California San Diego campus, La Jolla, CA, USA between June 20th and July 25th, 2023 (supplementary material, Figure S1). We refer to the ants collected from each location as a separate ‘collection’. The collected ants were kept in a temperature-regulated laboratory room and were fed with ant nectar (byFormica Sunburst Ant Nectar) and water *ad libitum*.

### Experimental apparatus and procedure

A custom-built experimental apparatus was used to record the body kinematics and arolium engagement in Argentine ants (Figure 1*a*). The apparatus consisted of a vertically mounted Frustrated Total Internal Reflection (FTIR) sensor, a 3D printed white bottomless tunnel (PLA), a Phantom VEO 410L high-speed camera (Vision Research, Wayne, NJ, USA), and two plastic containers (Figure 1*a*,*b*).

**Figure 1:**
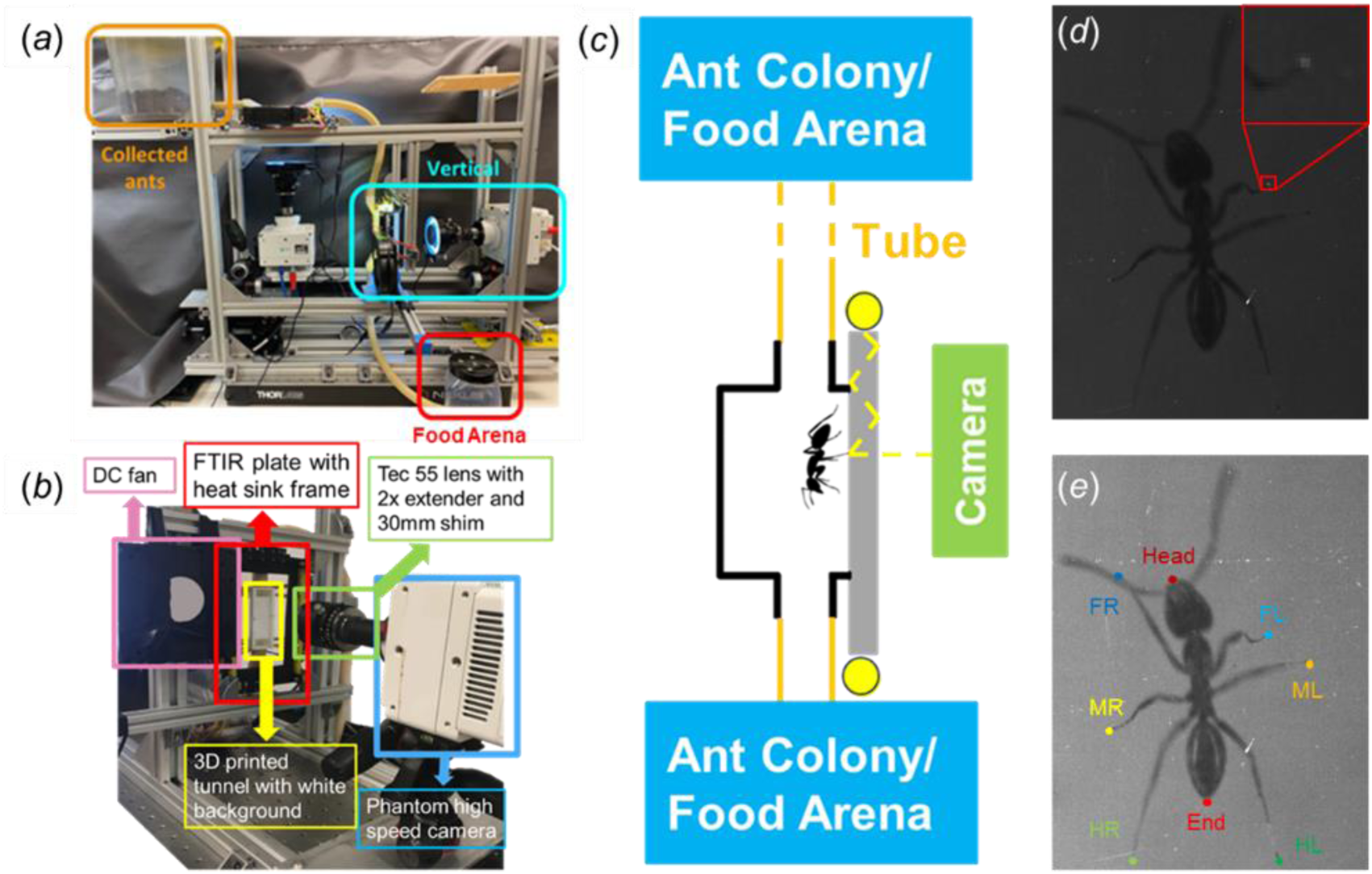
Experimental set-up. (***a***) Overall experimental set-up. (***b***) Detailed set-up for the vertical segment. (***c***) Diagram of the experiment. (***d***) Raw image of an ant during upward climbing. In this figure, FL and MR feet tips were illuminated indicating the engagement of arolia on these 2 legs. The illuminated FL arolium engagement is magnified and displayed in the red box. (***e***) Ant image with body parts labeled using DeepLabCut. The image shown is a contrast-adjusted version of (***d***), with contrast modified solely for visualization purposes. For tarsal tip labels, “F”, “M”, and “H” indicate front, middle, hind foot, respectively; “L” and “R” indicate left and right.

The FTIR sensor is a borosilicate glass plate (152.4mm × 101.6mm x 9.5mm; McMaster-Carr, Elmhurst, IL, USA) surrounded by a 13mm wide 24V LED light strip (6000K±300K, Daylight White; JOYLIT, Shenzhen, China). When light travels through the glass plate, light rays that are at an angle below the angle of refraction with respect to the plate surface will be reflected internally, as the glass plate has a higher index of reflection than its surrounding air. However, when an object such as the ant’s arolium is in nanoscale proximity to the outer plate surface, light can be transmitted through the surface (Figure 1*c*). The transmitted light from close contact points is then scattered out of the glass plate and toward a light-sensitive camera, where it is recorded as an illuminated spot (Figure 1*d*). This technology has been used previously to analyze animal foot contact [16–18].

During the experiment, ants can freely climb on the FTIR sensor (the temperature in the ant climbing area is controlled in a range of 22∼34°C [19,20]; supplementary material, Figure S2). The high-speed camera was set in Image Based Auto-Trigger (IBAT) mode with 400 frame per second (fps) recording frequency. Once ants entered the field of view, the camera automatically captured videos, with each video approximately 618 frames long. The recording session for each collection lasted 24 hours at a minimum.

### Step selection

The open-source toolbox DeepLabCut (version 4.2.3) was used to track the six tarsal tips of the feet, the anterior end of the head (“Head”) and the posterior end of the gaster (“End”) for each frame (Figure 1e). The Center of Body (CoB) was the middle point between the “Head” and “End”. All videos were separated into steps (from touch-down to lift-off for a hind limb; see supplementary material, Method S1 for details on how steps were separated). For each step, the tripod coordination strength (TCS) was calculated as the ratio between the number of frames in which all three legs of a given tripod were simultaneously in swing phase and the number of frames in which at least one of these legs was in swing phase [21]. Steps with TCS > 0.5 and orientated vertically (±20 degrees from the gravity direction) were selected for further analyses. Additionally, within each step, only frames where the ant was in an alternating tripod gait were included in the further analyses. The normalized time for each frame was calculated according to its position in the whole step frame sequences with 0 and 1 indicating the beginning and end of a step, respectively.

### Arolium engagement calculation

We identified arolium engagement by comparing pixel brightness at the foot with respect to the surrounding background noise (See supplementary material, Method S2 for complete description). Specifically, we used two metrics: 1) a binary representation of whether arolium are engaged/not engaged which is computed by thresholding the arolium light intensity, and 2) a measure of arolium engagement intensity which is computed by summing all pixels that exceed the noise threshold.

We verified that the illuminated contacts originated specifically from the arolium and not from other leg structures using a supplementary experiment (See supplementary material, Method S3 and Figure S3). Because the fine contact details of the arolium are below our optical resolution, we used the integrated FTIR light flux (summed pixel values) from each illuminated contact as a relative measure of total contact area, which we defined as engagement intensity. This interpretation is based on the physical principle of frustrated total internal reflection: a larger area of optical contact scatters more light, resulting in a higher total pixel intensity.

### 3D tripod quasi-static force model

A quasi-static tripod model of climbing was constructed as a point mass with three weightless limbs representing the ant’s center of mass connected with the 3 stance phase limbs. The center of mass (CoM) is located a distance ℎ from the surface in the z direction, and the 3 stance phase limbs are contacting the climbing surface. The mass point is subjected to gravity (*m**g***), and 3-dimensional foot contacting forces (***F***_1_, ***F***_2_, ***F***_3_) act on the tip of all stance-phase limbs (supplementary material, Figure S4). To maintain the quasi-static force balance, the sum of gravitational (*m**g***) and foot contact forces (***F***_1_, ***F***_2_, ***F***_3_) must be zero in all three directions, and the sum of torques about the center of mass due to the foot contact forces (***τ***_1_, ***τ***_2_, ***τ***_3_) must be zero for all three axes. These requirements allow us to get the analytic solution for the foot contacting forces in the normal directions (*F*_1*z*_*, F*_2*z*_*, F*_3*z*_; See supplementary material, Method S4 for details):

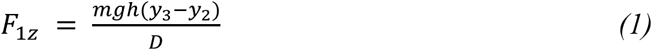

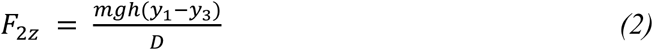

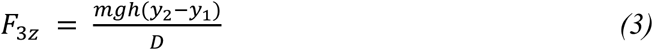

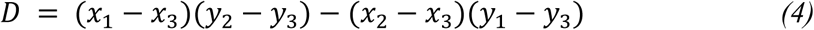

Where *F*_1*z*_*, F*_2*z*_*, F*_3*z*_ represent the signed scalar normal components of the foot contact forces. *mg* represents the magnitude of the gravitational force acting on the object, ℎ is a positive scalar representing the distance between the point mass and the contacting surface. *x*_1_*, x*_2_*, x*_3_ and *y*_1_*, y*_2_*, y*_3_ correspond to the x and y signed scalar components of the foot contact points relative to the center of mass (CoM). When calculating the required normal force, the measured center of body (CoB) position was used as an estimate of the CoM position, and the distance ℎ was assumed to remain constant during each step. The potential influence of the variations in *h* on the force predictions is assessed in the Discussion. As the locations of the contacting feet do not change relative to each other during each stance phase, the time dependencies of the foot positions (*x_i_*, *y_i_*) caused by the movement of the center of mass compensate in equations (1–3). This leads to step-wise time-invariant normal forces with positive values indicating compression forces and negative values indicating adhesion forces.

### Acceleration calculation

The instantaneous speed at each frame was determined by first calculating the displacement between the center of body (CoB) positions at the previous and next frames. Then the component of this displacement that aligned with the fore-aft body axis was taken, and divided by twice the frame duration (central difference method). Acceleration was obtained by applying the same central difference method to the previously computed instantaneous speed. The maximum magnitude of acceleration for each step was defined as the largest absolute value of all frame-wise accelerations.

### Statistical analysis

All statistical analyses were performed in R (v.4.2.3). Due to repeated measurements from different collections of ants from distinct locations and large differences in sample size across collections, we treated each collection as our experimental unit and used a two-stage method for all statistical analyses [22]. The two-stage method proceeded as follows: 1) a separate binomial (linear) regression model was fitted to each collection to obtain the estimate and its standard error per collection; 2) the collection-specific results were synthesized into an overall result using random-effects meta-analyses (metafor package, [23]).

In the first stage, binomial regression models were used to obtain the estimates and standard errors for the arolium engagement probability across foot positions, stance types, and normalized time. Linear regression models were used to estimate the relationship between the log-transformed engagement intensity ratio and the log-transformed adhesion force ratio, and the temporal variation of acceleration. Second order polynomial regression models were used to estimate the quadratic coefficients (*a*) and the linear coefficients (*b*) for the temporal variation of engagement intensity. Vertices (peaks) were calculated as 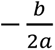. Standard errors for vertices were obtained using the delta method.

In the second stage, univariate meta-analyses were performed to obtain overall estimates and to test our hypotheses. To test whether outcomes varied with direction (up vs. down) or stance type (narrow vs. wide), we calculated the corresponding collection-specific differences and synthesized each set of differences using meta-analysis. In our study, we chose 0.05 as the level for statistical significance (α). To account for multiple testing, we applied Holm-Bonferroni correction [24] to p-values from tests of whether (1) middle arolium engagement probability varied with direction or stance type, (2) engagement probability varied over time, and (3) engagement intensity showed an asymmetric temporal pattern.

## Results

### 1. Stance type and arolium engagement probability

We first checked if the arolia that are required to generate adhesion forces exhibited arolium engagement (as indicated by FTIR illumination). According to the quasi-static model, the uppermost foot should always generate adhesion force, while the lowest arolium should always generate compression force, regardless of climbing direction. The required normal force direction for middle arolium is also direction-independent but depends on whether an ant is in wide stance or narrow stance (Figure 2*a*). Narrow stance is when the lowest foot is laterally closer to the center of the body than the uppermost foot. Wide stance is when the lowest foot is laterally further from the center of the body than the uppermost foot. If an ant is in a narrow stance, their middle leg would be required to generate adhesion force, while if an ant is in a wide stance, their middle leg would be required to compress. During upward climbing, ants exhibited a wide stance within 79.1% of observations, while during downward climbing, they primarily exhibited a narrow stance in 76.6% of observations (Figure 2*b*).

**Figure 2:**
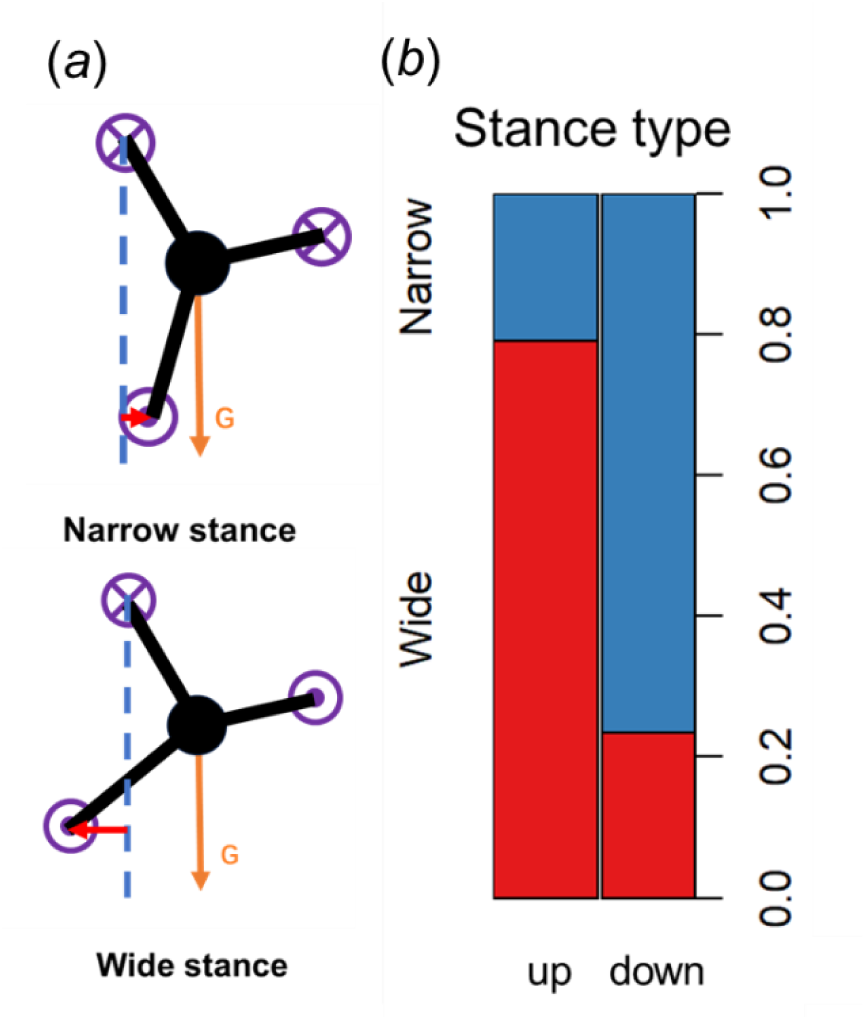
Stance types and model-predicted normal forces. (***a***) Diagram of narrow stance and wide stance. Black symbols are the tripod models in the dorsal plane. Orange arrows labeled with “G” represent gravity. Blue dashed lines and red arrows show the relative lateral position between the uppermost foot and the lowest foot. The purple symbols indicate the required normal force direction for each foot according to the tripod force model. ⊗ denotes an adhesion force, while ⊙ denotes a compression force. (***b***) Stance type probability in upward and downward climbing. Red bars represent the probability of a wide stance, while the blue bars show the probability of a narrow stance.

The measured engagement probabilities of uppermost and lowest arolia showed significant differences between upward and downward climbing (Figure 3*a*). During upward climbing, the uppermost (front) foot showed arolium engagement probability of 96% [94%, 97.3%], which was significantly higher (*z* = 17.9, *p* < 0.05, *k* = 8) than the uppermost (hind) arolium engagement probabilities (53.3% [40.1%, 66.1%]) during downward climbing. Conversely, the lowest (hind) arolium had significantly lower engagement probability (*z* = -10.88, *p* < 0.05, *k* = 8) during upward climbing (1.2% [0.6%, 2.3%]) than during downward climbing (15.3% [8.9%, 25%]). During upward climbing, the engagement status of uppermost and lowest arolia matched the required normal forces predicted by the quasi-static model in over 90% cases, while during downward climbing, less than 70% matched.

**Figure 3:**
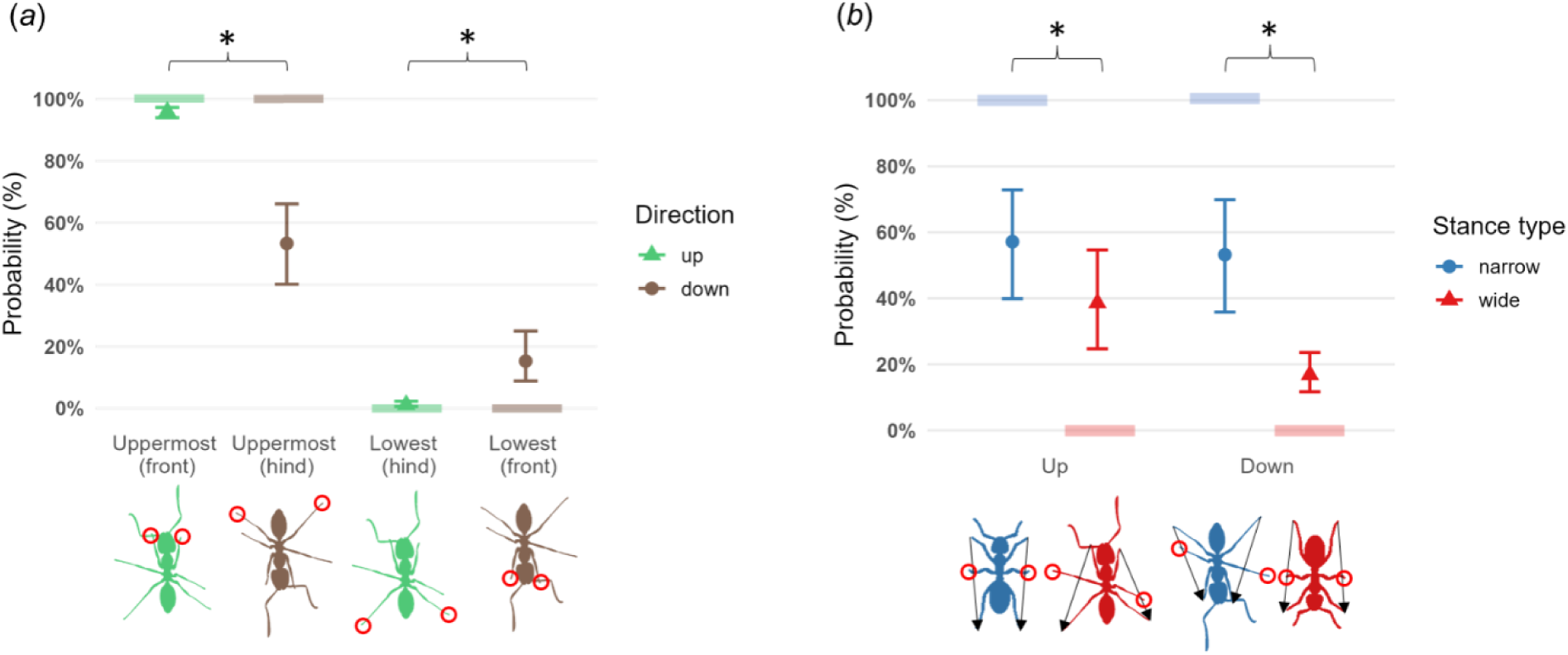
Arolium engagement probability. (***a***) Engagement probabilities for uppermost and lowest arolia. During upward climbing, the uppermost foot is the front foot and the lowest foot is the hind foot. During downward climbing, this relationship reverses: the uppermost foot is the hind foot and the lowest foot is the front foot. (***b***) Engagement probability for middle arolium. For both (***a***) and (***b***), points represent the pooled probability estimates across all collections and the error bars show their 95% confidence intervals. The semi-transparent lines at probabilities 0% and 100% indicate the predictions from the quasi-static tripod model. The asterisks (*) indicate significant differences.

The measured engagement probabilities of middle arolium did not differ significantly between upward (narrow: 57.1% [39.9%, 72.8%]; wide: 38.6% [24.7%, 54.6%]) and downward climbing (narrow: 53.2% [35.8%, 69.9%]; wide: 16.8% [11.7%, 23.6%]) for both stance types (narrow: *z* = 0.92, Holm-Bonferroni adjusted *p* = 0.36, *k* = 8; wide: *z* = 2.2, Holm-Bonferroni adjusted *p* = 0.056, *k* = 8). The middle arolium exhibited significant differences between wide stance and narrow stance when ants climbed in both directions (up: *z* = -3.08, Holm-Bonferroni adjusted *p* < 0.05, *k* = 8; down: *z* = -7.89, Holm-Bonferroni adjusted *p < 0.05, k = 8*; Figure 3*b*). The probability that the engagement status matched the required normal forces ranged from 53.2% to 83.2% for middle arolium. Forest plots showing variances across collections are provided in supplementary material, Figures S5 and S6.

### 2. Relationship between arolium engagement intensity ratio and required adhesion force ratio

For steps where the quasi-static model predicted that both the uppermost and the middle feet (front and middle during upward climbing; hind and middle during downward climbing) needed to generate adhesion forces (*F*_1*z*_, *F*_2*z*_) and the observed arolium engagement intensities (*I*_1_, *I*_2_) were non-zero, we tested whether the leg requiring higher adhesion force showed more intense arolium engagement. Specifically, we analyzed the relationship between the arolium engagement intensity ratio (*I*_1_/*I*_2_) and the required adhesion force ratio (*F*_1*z*_/*F*_2*z*_) of the two limbs during upward and downward climbing (Figure 4*a*). The engagement intensity ratios increased with the required adhesion force ratios for both climbing directions. During upward climbing, the engagement intensity ratio (*I*_1_/*I*_2_) scaled as (*F*_1*z*_/*F*_2*z*_)^0.38^ (95% CI: 0.30-0.46, *z* = 9.45, *p* < 0.05, *k* = 8, Figure 4*b*), and during downward climbing, the engagement intensity ratio (*I*_1_/*I*_2_) scaled as (*F*_1*z*_/*F*_2*z*_)^0.3^ (95% CI: 0.16-0.44, *z* = 4.13, *p* < 0.05, *k* = 8, Figure 4*c*). Although the overall scaling exponent was positive for downward climbing, individual scaling exponents for collections C0620, C0627 and C0703 were negative. Forest plots showing scaling exponents for individual collections are provided in supplementary material, Figure S7.

**Figure 4:**
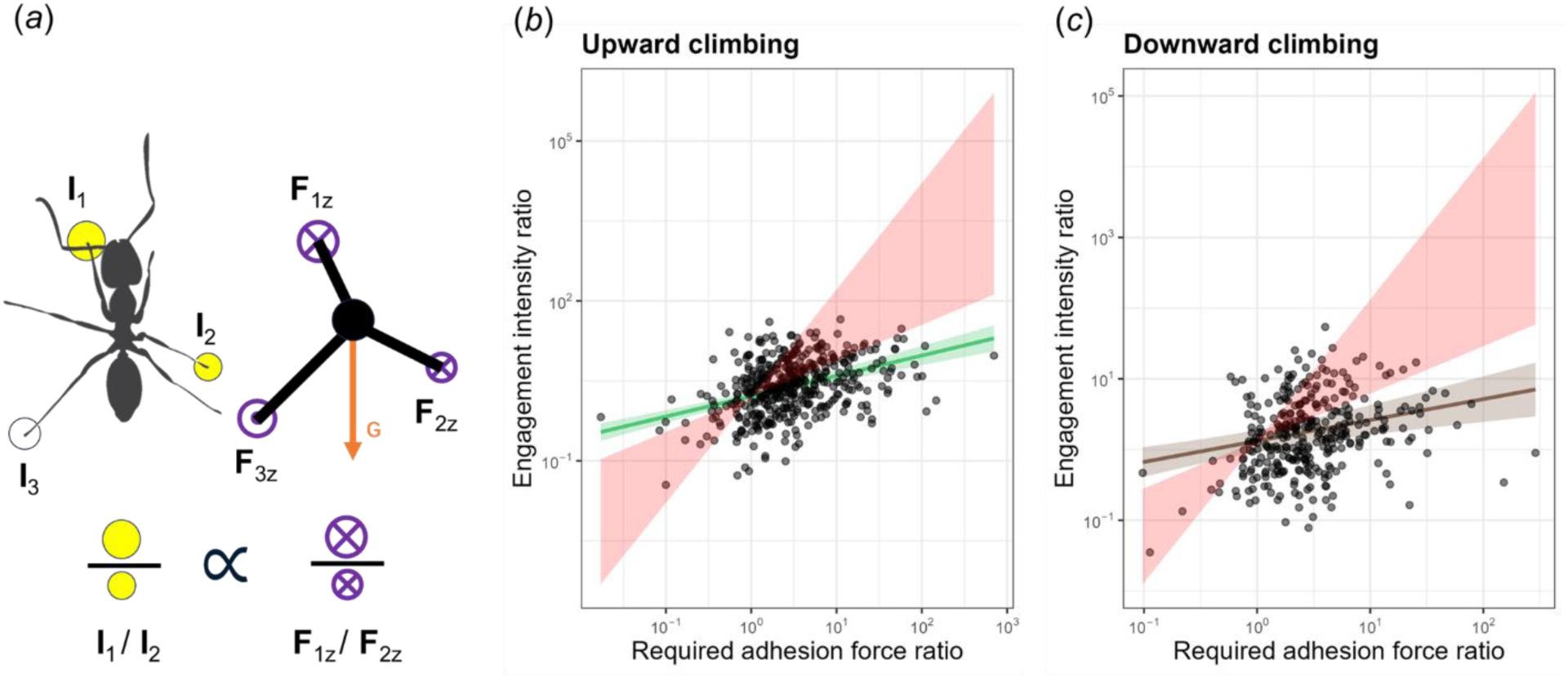
Arolium engagement intensity ratio and required adhesion force ratio. (***a***) Diagram showing the calculation of arolium engagement intensity and adhesion force ratios. The orange arrow indicates the gravity direction. Circles labeled as I_1,2,3_ represent the arolium engagement intensities with the white one indicating the zero intensity and yellow ones indicating non-zero intensities. The size of each yellow circle represents the magnitude of the arolium engagement intensity. ⊗ labeled as *F*_1*z*,2*z*_ denotes the required adhesion force, while ⊙ labeled as *F*_3*z*_ denotes the required compression force. The size of ⊗ represents the magnitude of the required adhesion force predicted by the tripod model. We hypothesize that the engagement intensity ratio (*I*_1_/*I*_2_) between two different arolia should be positively related to the ratio of their required adhesion forces (*F*_1*z*_/*F*_2*z*_). (***b***,***c***) Relationship between arolium engagement intensity ratio and required adhesion force ratio during climbing up (***b***) and climbing down (***c***). For both (***b***) and (***c***), the fitted line shows the overall power-law relationship synthesized across collections, with the shaded area indicating the 95% confidence interval (green for up, brown for down). The red shaded area represents the expected range of the power-law relationship between real contact area and adhesion force. Note that both axes are on logarithmic scales (log-log plot).

However, the scaling exponents in both climbing directions were significantly lower than the expected range (0.67–2), as the entire 95% CIs fell below 0.67 (Figure 4*b*,*c*). The expected range was estimated based on the pull-off adhesion forces generated by individual arolium and by whole-animal [25,26]. Adhesion stress was found increasing with shear force acting on the arolium. From a case with zero shear force to a case with shear force equal to body weight, the adhesion force (*F_a_*) generated by arolium was expected to vary from *m*^1/3^ to *m*^1^ (*m* is the body mass). Under isometric scaling (geometric similarity), arolium area (*A*) is predicted to scale with body mass as *A∼m^2/3^*, so we expected the observed scaling exponent between the engagement intensity ratio (a proxy for real contact area) and the required adhesion force ratio to fall within the range of 0.67 to 2 (*A*∼*F_a_*^0.67−2^).

### 3. Arolium engagement with respect to time

We next investigated whether the engagement status of each arolium stayed the same during a stance by studying: 1) the probability of arolium engagement over time, and 2) the intensity of arolium engagement over time. When it come to the engagement probability during downward climbing, temporal variation was significant for lowest (*z* = 15.99, Holm-Bonferroni adjusted *p* < 0.05, *k* = 8) and uppermost arolia (*z* = -4.29, Holm-Bonferroni adjusted *p* < 0.05, *k* = 8), but not for middle arolium (narrow stance: *z* = -1.47, Holm-Bonferroni adjusted *p* = 0.57, *k* = 8; wide stance: *z* = 0.7, Holm-Bonferroni adjusted *p* = 1, *k* = 8, Figure 5*a*). During upward climbing, none of the arolia showed significant temporal variation in engagement probability (Holm-Bonferroni adjusted *p* > 0.05 for all arolia). When Argentine ants climbed down, the arolium engagement log-odds for uppermost arolium increased (pooled slope estimate: 2.26 [1.98, 2.54], Figure 5*b*), and the log-odds for lowest arolium decreased (pooled slope estimate: -1.29 [-1.88, - 0.7], Figure 5*c*) over the normalized time.

**Figure 5:**
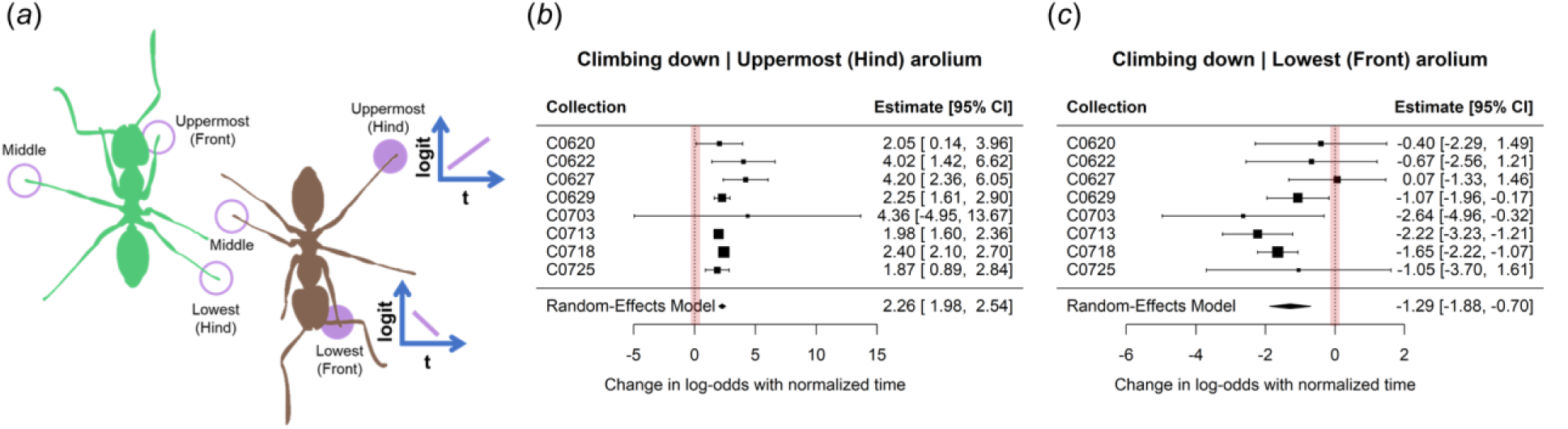
Temporal variation of arolium engagement probability. (***a***) Results overview. The ant body in green represents the climbing up scenario, while the ant body in brown represents the climbing down scenario. Feet labeled with purple circles are those in the stance phase. Among them, feet marked with solid circles showed a time-dependent engagement probability. The purple lines in the blue coordinates indicate whether the probability increases or decreases with respect of time. (***b***,***c***) Plots showing estimated slopes and 95% confidence intervals for the relationship between the log-odds of arolium engagement and normalized time across 8 collections. Each point represents the estimated slope from a single collection, and the horizontal lines indicate the corresponding 95% confidence intervals. The black diamond below the collection values shows the pooled slope estimate and its 95% confidence interval of all collections, and the vertical dashed line with red shade at slope = 0 indicates no temporal variation of engagement probability. Data to the left of the vertical dashed line (negative values) indicate a decreasing trend in engagement probability, whereas data to the right (positive values) indicate an increasing trend.

We then studied the arolium engagement intensity over time for adhering feet. Since a stance begins and ends with no contact area, we expected the temporal pattern of engagement intensity would increase to a maximum and then decrease, following a peak-shaped second-order polynomial pattern. The middle arolium showed a relatively weaker second-order polynomial pattern: more than three collections had adjusted *R^2^* values below 0.1, indicating substantial variance not explained by the polynomial model. For uppermost and lowest arolia, engagement intensities versus time across all collections showed a peak-shaped pattern. The only exception was the lowest (hind) arolium during upward climbing (supplement materials, Figures S8). Thus, we focus the remaining analysis on the uppermost arolium during upward climbing and uppermost and lowest arolia during downward climbing.

The arolium engagement intensities showed asymmetric temporal trend, indicated by the vertices (timestamp with peak engagement intensity) significantly different from the mid-stance (0.5).

During upward climbing, the engagement intensity of the uppermost (front) arolium rose briefly to an early peak and then mostly decreased for the rest of the step, indicated by a vertex significantly earlier than mid-stance (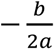 = 0.2, 95% CI: [0.12, 0.28], *z* = -7.47, Holm-Bonferroni adjusted *p* < 0.05, *k* = 8, Figure 6*a*). In contrast, during downward climbing, the arolium engagement intensity of the uppermost (hind) arolium mostly increased, with a brief decline toward the end of the step. This trend was indicated by a vertex significantly later than mid-stance (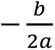 = 0.69, 95% CI: [0.66, 0.72], *z* = 13, Holm-Bonferroni adjusted *p* < 0.05, *k* = 8, Figure 6*b*). The arolium engagement intensity of the lowest (front) arolium mostly decreased after a brief incline, indicated by a vertex significantly earlier than mid-stance (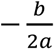 = 0.22, 95% CI: [0.08, 0.36], *z* = -4, Holm-Bonferroni adjusted *p* < 0.05, *k* = 8, Figure 6). Forest plots showing vertices for individual collections are provided in supplementary material, Figure S9.

**Figure 6:**
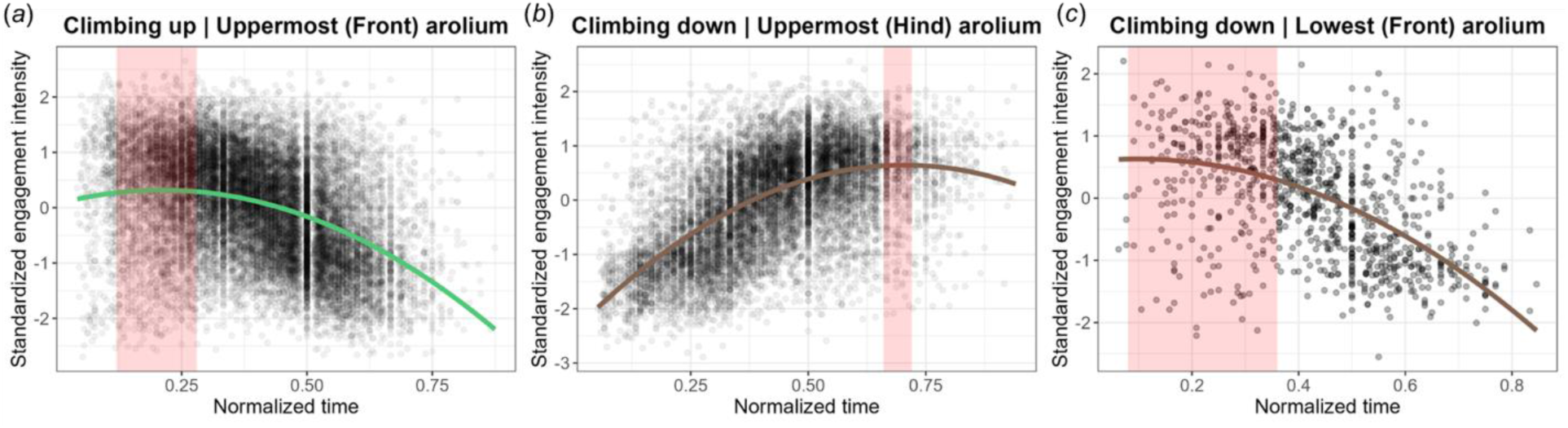
Engagement intensity over normalized time for uppermost and lowest arolia. (***a***-***c***) The intensity was standardized by subtracting its mean and dividing by its standard deviation within each step. Gray points are the data across all collections. The fitted curve (green for up, brown for down) is based on the pooled coefficients obtained from the meta-analysis across collections. The red shaded area indicates the 95% confidence interval for the pooled vertex.

## Discussion

Small insects like Argentine ants climb smooth surfaces through the use of adhesive pads on their feet called arolium. In this work, we tested whether Argentine ants distribute their arolium engagement on different legs according to the required normal forces predicted by a quasi-static model. Our results indicated that certain aspects of arolium engagement aligned with the normal force requirements. For instance, during upward climbing, the uppermost (front) arolium always engaged and the lowest (hind) arolium mostly disengaged, closely matching the quasi-static normal force requirement. In addition, middle arolium engagement status showed dependence on the stance-type but not on the climbing direction. However, the discrepancies between the measured arolium engagement and the quasi-static normal force requirement were more pronounced. For example, during downward climbing, the arolium engagement pattern was not symmetric to that during upward climbing, and the match rate with normal force requirement was less than 70%. Moreover, some arolia showed temporal variation significantly different from the baseline expectation.

### Arolia engagement probability differed significantly when ants climbed in different directions

Contrary to the direction-independent force requirement (uppermost arolium generates adhesion force, while lowest generates compression force), climbing direction influenced the arolium engagement probability. During downward climbing, the uppermost (hind) arolium engagement probability was significantly lower, and the lowest (front) arolium engagement probability was significantly higher (Figure 5), compared to the results during upward climbing. There are multiple potential factors for this difference.

The first potential factor is the difference in body acceleration when Argentine ants climbed in different directions. Ants achieved significantly greater peak acceleration magnitudes (*z* = 10.61, *p* < 0.05, *k* = 8; Figure 7*a*) during downward climbing (0.23g [0.19,0.26]) than during upward climbing (0.15g [0.12,0.17]). Higher acceleration may make downward climbing less predictable by our quasi-static model compared to upward climbing and can potentially cause bigger variances in engagement probabilities as seen in Figure 3*a*.

**Figure 7:**
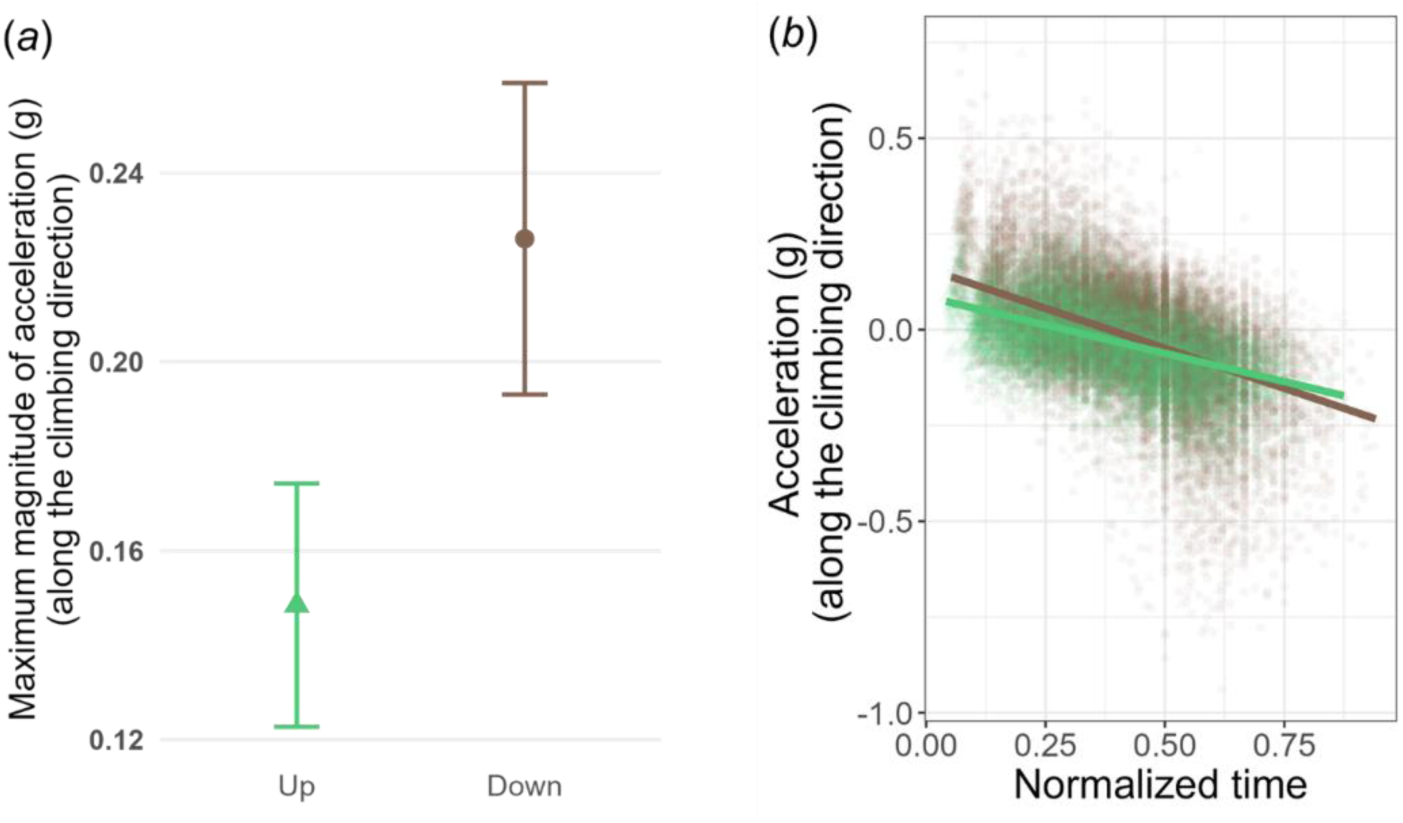
Acceleration along the climbing direction. (***a***) Maximum magnitude of acceleration. Points represent the pooled estimates across all collections and the error bars show their 95% confidence intervals. (***b***) Temporal variation of acceleration. Green points are the climbing up data across all collections, and the green straight line is the fitted linear regression line. The brown line and brown points represent the results for climbing down. g is 9.8m/s^2^.

Furthermore, the non-isotropic body mass distribution may contribute to the observed engagement probability differences between upward and downward climbing. The quasi-static model simplified the ant body to a mass point, thereby ignoring the uneven distribution of body mass in Argentine ants. Changing mass distribution can alter limb forces, as a previous study in trotting dogs showed that changing load distribution altered individual limb mechanics and contact timing [27]. This suggests that the model’s simplification may partly explain why the required normal force is direction-independent, while the engagement probability is not.

Lastly, we assumed foot-only substrate contact in our study. However, studies on Argentine ant trailing behavior have shown that other body parts can contact the substrate [3,28,29]. For example, when ants walked upside-down on a glass slide, their gasters were observed touching the glass [3]. When ants walked on an inclined tube, pheromones on the substrate were deposited from the gaster, and ants were observed touching the substrate with their antennae [29].

Therefore, other body parts may occasionally contact the substrate during vertical climbing. If antennae, mandibles, or the gaster contact the substrate and influence climbing differently depending on direction, such contact could potentially contribute to direction-dependent arolium engagement.

### Middle arolium exhibited intermediate engagement probability with high variance

Consistent with the quasi-static normal force requirement, the engagement probability of middle arolium was direction-independent but varied with stance type. However, unlike the required normal forces, its engagement probability was never close to 100% for a narrow stance case or near 0% for a wide stance case and its engagement probability always varied over a wide range.

The intermediate and inconsistent engagement of the middle arolium may result from a combination of its low normal force requirement and the absence of a consistent force pulling toward or pushing against the body. The low normal force requirement can be derived from the quasi-static model. Within a stance of the alternating tripod gait, the lateral distance between the middle and the lowest feet is often greater than that between the front and the lowest feet, which, according to the quasi-static model, results in a lower required adhesion force for the middle arolium.

Previous studies have shown that the extension of arolium can be controlled passively and the passive mechanism is direction-dependent [6,11,30]. The arolium extends if the corresponding leg is pulled toward the body, while the arolium folds when the leg is pushed against the body. During vertical climbing, compared to the front or the hind leg, the middle leg is typically oriented more perpendicularly to the direction of gravity, resulting in reduced passive shear force caused by gravity along its axis.

Furthermore, lateral body oscillation was observed in our climbing ants. Such oscillation may require pulling or pushing forces along the middle leg’s direction, as reported in a previous study on wood ants running [31]. Since the required normal force and the consistent pulling or pushing force caused by gravity are small, the inconsistent pushing and pulling forces required by lateral body oscillation may dominate the engagement status of the middle arolium. Additionally, a low required normal force necessitates a small engagement intensity, which may be difficult to detect accurately.

### Association between engagement intensity and required adhesion force was weak and deviated from the expected range

When it comes to the overall relationship between the engagement intensity ratio and the required adhesion force ratio, the scaling exponents were significantly positive for both climbing directions. However, some collections during downward climbing had negative scaling exponents. In addition, the data points were widely scattered around the fitted relationship, suggesting a large portion of the total variance in engagement intensity ratio remained unexplained.

One potential explanation for this is that, during free climbing, ants engage their arolia across a broad range of contact areas that are all sufficient to meet both adhesion force and speed demands. This points to a functional plateau rather than a finely tuned optimum. Supporting this idea, ants exhibited high acceleration/deceleration during some fast steps (supplementary material, Figure S10). This likely requires both increased adhesive force to prevent detachment, and short step duration to maintain fast speed. If ants can achieve both strong adhesion force and rapid movement under such conditions, during slower steps—when both adhesion force and speed demands are lower—a range of intermediate contact areas would be sufficient. Thus, instead of optimizing contact area according to required adhesion force for every step, ants may rely on a robust adhesive system that performs effectively across a range of engagement levels. A previous study on geckos also found the same phenomenon, as they revealed setae might make contact without generating a significant normal force [32].

In addition to the large amount of unexplained variance, the measured correlation parameter was out of the expected adhesion range. This may result from a nonlinear relationship between light flux and real contact area. Although the light flux increases with real contact area in FTIR studies, a previous work with a cylindrical puck pressed against an FTIR sensor suggests that deformation of the contact material can alter scattering intensity [33].

### Arolia exhibited time-dependent engagement probability and intensity during each step

The quasi-static model predicts that throughout each climbing step, ants’ foot normal forces do not change over time, even as their center of mass (CoM) moves relative to the supporting legs. Contrary to the predicted time-independence of the required adhesion force, in experiment we observed that the uppermost and lowest arolia during downward climbing exhibited time-dependent engagement probabilities. In addition, the uppermost arolium when climbing in both direction and lowest arolium during downward climbing showed significant asymmetric temporal changes in engagement intensity.

Time dependence of engagement intensity may come from the inherent cyclic nature of a stance (initiate engagement, adhere, disengage), or be related to locomotion mode. To assess the source of this time dependence, we compared the temporal changes in engagement intensity to previously published data on temporal variations in contact area of Asian Weaver ants (*Oecophylla smaragdina*) during upside-down walking [10] (supplementary material, Figure S11). The vertices (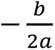 ) for the upside-down data were much closer to mid-stance (0.51 for middle arolium and 0.56 for hind arolium), suggesting more symmetrical engagement patterns compared to the trends we measured during climbing. These comparisons supported that during climbing, the asymmetric temporal variations in engagement intensity were not because of the cyclical nature of the engagement and disengagement processes but non-inherent changes related to climbing.

Among cases with time-dependent engagement, the uppermost arolium during downward climbing displayed an increasing trend in both engagement probability and intensity, whereas all other cases showed decreasing trends. One reason proposed by us to explain the time-dependent engagement for uppermost and lowest arolia is the non-negligible body acceleration during climbing.

During each step, Argentine ants accelerated first and then decelerated (Figure 10B; up: pooled slope = -0.3 [-0.39, -0.2], *z* = -6.15, *p* < 0.05, *k* = 8; down: pooled slope = -0.42 [-0.56, -0.28], *z* = -5.94, *p* < 0.05, *k* = 8). This temporal pattern of acceleration aligned with the temporal trend of the uppermost arolium engagement for both upward and downward climbing (Figure 5,6). In the static case, the normal force requirement (Equation 3-6) is influenced by gravity alone. However, during locomotion, the body experiences both gravitational force (mg) and inertial forces due to acceleration (ma). During upward climbing, the ant’s body first experiences a net upward force to overcome gravity and accelerate, then a net downward force as gravity assists in deceleration. The normal force requirement must therefore adjust to reflect this changing net load, necessitating the adhesion force on the top arolium to be stronger at the start and then decrease, aligning with the arolium engagement intensity trend in Figure 6*a*. Conversely, during downward climbing, ants use gravity to accelerate initially and later need more upward force to decelerate, causing the adhesion force on the top arolium to be weaker at first and increase over time, matching the arolium engagement trend in Figures 5*b* and 6*b*.

This acceleration trend during downward climbing also requires the lowest arolium to gradually increase the compressing force. As ants often do not adhere their arolium when the corresponding leg presses onto the surface [9], we expect the lowest arolium engagement intensity to change in the opposite direction of the compressing force requirement. The measured decreasing trend (Figures 5*c* and 6*c*) matched our expectation. Thus, we hypothesize that body accelerations are important in ant climbing dynamics, which can be one explanation for why the quasi-static force requirement fails to predict the observed temporal engagement trends.

Another possible explanation is that the distance between the ant’s center of mass and the contacting surface may vary over time. Previous studies on level walking in *Formica pratensis, Cataglyphis fortis* [34] and *Formica polyctena* [35] and slope traversing in *Formica pratensis* and *Cataglyphis fortis* [34] reported small body height oscillations, with height variance below 10% of body height in most cases, supporting the validity of our constant CoM-to-substrate distance assumption in calculating normal force requirement. However, it is unknown whether CoM-to-substrate distance oscillations during vertical climbing remain similarly small.

Therefore, future studies should incorporate three-dimensional body kinematics to account for varying CoM-to-substrate distance when calculating the normal force requirement.

### Comparison with previous studies suggests some findings may apply beyond Argentine ants

Consistent with the asymmetric arolium engagement pattern, a study on *Cataglyphis fortis* climbing slopes reported asymmetric leg impulses between ascending and descending. When *Cataglyphis fortis* walked on a 60° upslope, the front (uppermost) legs mostly generated compression impulses in the normal direction, while on a 60° downslope, the hind (uppermost) legs mostly generated adhesion impulse [36].

During level walking, *Formica polyctena* displayed acceleration ranging from approximately -2 m/s² to 1.5 m/s² within a single step [35], which matches the magnitude we measured in Argentine ants during vertical climbing (up: from -1.69 m/s^2^ to 0.72 m/s²; down: from -2.28 m/s^2^ to 1.35 m/s²). This agreement suggests that arolium engagement in other ant species may also be impacted by within-step acceleration.

Moreover, such non-negligible within-step acceleration was not only observed in ants, but also in other insects. During rapid vertical climbing, cockroaches showed fore-aft forces oscillating around the body weight. The peak fore-aft acceleratory forces generated by cockroaches were around 1.7 times the body weight, while the minimum values were around 0.26 times the body weight [37], indicating an acceleration range from -6.27 m/s^2^ to 6.86 m/s^2^ within a stride. Based on Figure 4 in [38], within a fast step, the speed change of *Drosophila* could go up to around 60% of its average speed, indicating a non-negligible acceleration.

## Conclusion

In conclusion, the measured arolium engagement of Argentine ants showed asymmetric pattern between upward and downward climbing, and changed over time within a single step. These characteristics could not be explained by the normal force requirement of a quasi-static model during vertical climbing. This disagreement may arise from several possible factors including: (1) non-negligible within-step body accelerations; (2) anatomical complexities, including unevenly distributed body mass that violate the simplified point-mass assumption of the model; (3) passive mechanical coupling between leg forces and arolium extension and (4) the ants’ flexibility to select contact areas within a broad range that appears sufficient to meet both adhesion and speed demands. Future studies incorporating simultaneous force measurements across feet may help determine the arrangement and control of arolium engagement in rapid climbing.

## Acknowledgements

We thank Dr. David Holway, Dr. David Labonte and Dr. Nicholas Boechler for their valuable suggestions on this project. We also thank Dr. Glenna Clifton, and all members of the Gravish lab (2022–2025) for their advice and help in implementing the experiments. We acknowledge funding from NSF-CAREER #2048235.

## Data availability

Datasets and R code for statistical analyses are available in the Dryad Digital Repository: https://doi.org/10.5061/dryad.cz8w9gjkd. Raw videos are available from the corresponding author upon reasonable request.

## References

1. Mallis A. 1942 Half a century with the successful Argentine ant. Sci. Mon. 55, 536–545.

2. Choe DH, Villafuerte DB, Tsutsui ND. 2012 Trail pheromone of the Argentine ant, Linepithema humile (Mayr) (Hymenoptera: Formicidae). PLoS ONE 7, e45016. (doi:10.1371/journal.pone.0045016)

3. Reid CR, Latty T, Beekman M. 2012 Making a trail: informed Argentine ants lead colony to the best food by U-turning coupled with enhanced pheromone laying. Anim. Behav. 84, 1579–1587. (doi:10.1016/j.anbehav.2012.09.036)

4. Brechka P. 2024 Slippery surfaces and the biomechanics of climbing in Macaranga-Ant mutualisms. PhD thesis. Cambridge, UK: University of Cambridge.

5. Endlein T, Federle W. 2008 Walking on smooth or rough ground: passive control of pretarsal attachment in ants. J. Comp. Physiol. A 194, 49–60. (doi:10.1007/s00359-007-0287-x)

6. Federle W, Brainerd EL, McMahon TA, Hölldobler B. 2001 Biomechanics of the movable pretarsal adhesive organ in ants and bees. Proc. Natl. Acad. Sci. USA 98, 6215–6220. (doi:10.1073/pnas.111139298)

7. Gorb SN. 2007 Smooth attachment devices in insects: functional morphology and biomechanics. In Advances in Insect Physiology, vol. 34 (ed. J Casas), pp. 81–115. London: Elsevier. (doi:10.1016/S0065-2806(07)34002-2)

8. Federle W, Rohrseitz K, Hölldobler B. 2000 Attachment forces of ants measured with a centrifuge: better ‘wax-runners’ have a poorer attachment to a smooth surface. J. Exp. Biol. 203, 505–512. (doi:10.1242/jeb.203.3.505)

9. Endlein T, Federle W. 2015 On heels and toes: how ants climb with adhesive pads and tarsal friction hair arrays. PLoS ONE 10, e0141269. (doi:10.1371/journal.pone.0141269)

10. Federle W, Endlein T. 2004 Locomotion and adhesion: dynamic control of adhesive surface contact in ants. Arthropod Struct. Dev. 33, 67–75. (doi:10.1016/j.asd.2003.11.001)

11. Federle W, Riehle M, Curtis ASG, Full RJ. 2002 An integrative study of insect adhesion: mechanics and wet adhesion of pretarsal pads in ants. Integr. Comp. Biol. 42, 1100–1106. (doi:10.1093/icb/42.6.1100)

12. Autumn K, Hsieh ST, Dudek DM, Chen J, Chitaphan C, Full RJ. 2006 Dynamics of geckos running vertically. J. Exp. Biol. 209, 260–272. (doi:10.1242/jeb.01980)

13. Wang Z, Wang J, Ji A, Zhang Y, Dai Z. 2011 Behavior and dynamics of gecko’s locomotion: the effects of moving directions on a vertical surface. Chin. Sci. Bull. 56, 573–583. (doi:10.1007/s11434-010-4303-2)

14. Humeau A, Piñeirua M, Crassous J, Casas J. 2019 Locomotion of ants walking up slippery slopes of granular materials. Integr. Org. Biol. 1, obz020. (doi:10.1093/iob/obz020)

15. Seidl T, Wehner R. 2008 Walking on inclines: how do desert ants monitor slope and step length? Front. Zool. 5, 8. (doi:10.1186/1742-9994-5-8)

16. Gravish N, Wilkinson M, Sponberg S, Parness A, Esparza N, Soto D, Yamaguchi T, Broide M, Cutkosky M, Creton C, Autumn K. 2010 Rate-dependent frictional adhesion in natural and synthetic gecko setae. J. R. Soc. Interface 7, 259–269. (doi:10.1098/rsif.2009.0133)

17. Mendes CS, Bartos I, Akay T, Márka S, Mann RS. 2013 Quantification of gait parameters in freely walking wild type and sensory deprived Drosophila melanogaster. eLife 2, e00231. (doi:10.7554/eLife.00231)

18. Wang X, Wang W, Tang Y, Wang H, Zhang L, Wang J. 2021 Apparatus and methods for mouse behavior recognition on foot contact features. Knowl. Based Syst. 227, 107088. (doi:10.1016/j.knosys.2021.107088)

19. Holway DA, Suarez AV, Case TJ. 2002 Role of abiotic factors in governing susceptibility to invasion: a test with Argentine ants. Ecology 83, 1610–1619. (doi:10.1890/0012-9658(2002)083[1610:ROAFIG]2.0.CO;2)

20. Human KG, Weiss S, Weiss A, Sandler B, Gordon DM. 1998 Effects of abiotic factors on the distribution and activity of the invasive Argentine ant (Hymenoptera: Formicidae). Environ. Entomol. 27, 822–833. (doi:10.1093/ee/27.4.822)

21. Wosnitza A, Bockemühl T, Dübbert M, Scholz H, Büschges A. 2013 Inter-leg coordination in the control of walking speed in Drosophila. J. Exp. Biol. 216, 480–491. (doi:10.1242/jeb.078139)

22. Gerow K, Bozeman B, Grossman GD. 2021 Response feature analysis for repeated measures in ecological research. Bull. Ecol. Soc. Am. 102, e01866. (doi:10.1002/bes2.1866)

23. Viechtbauer W. 2010 Conducting meta-analyses in R with the metafor package. J. Stat. Softw. 36, 1–48. (doi:10.18637/jss.v036.i03)

24. Holm S. 1979 A simple sequentially rejective multiple test procedure. Scand. J. Stat. 6, 65–70.

25. Labonte D, Struecker M-Y, Birn-Jeffery AV, Federle W. 2019 Shear-sensitive adhesion enables size-independent adhesive performance in stick insects. Proc. R. Soc. B 286, 20191327. (doi:10.1098/rspb.2019.1327)

26. Labonte D, Federle W. 2015 Scaling and biomechanics of surface attachment in climbing animals. Phil. Trans. R. Soc. B 370, 20140027. (doi:10.1098/rstb.2014.0027)

27. Lee DV, Stakebake EF, Walter RM, Carrier DR. 2004 Effects of mass distribution on the mechanics of level trotting in dogs. J. Exp. Biol. 207, 1715–1728. (doi:10.1242/jeb.00947)

28. Aron S, Pasteels JM, Deneubourg JL. 1989 Trail-laying behaviour during exploratory recruitment in the Argentine ant, Iridomyrmex humilis (Mayr). Biol. Behav. 14, 207–217.

29. Robertson PL, Dudzinski ML, Orton CJ. 1980 Exocrine gland involvement in trailing behaviour in the Argentine ant (Formicidae: Dolichoderinae). Anim. Behav. 28, 1255–1273. (doi:10.1016/S0003-3472(80)80114-X)

30. Endlein T, Federle W. 2013 Rapid preflexes in smooth adhesive pads of insects prevent sudden detachment. Proc. R. Soc. B 280, 20122868. (doi:10.1098/rspb.2012.2868)

31. Reinhardt L, Weihmann T, Blickhan R. 2009 Dynamics and kinematics of ant locomotion: do wood ants climb on level surfaces? J. Exp. Biol. 212, 2426–2435. (doi:10.1242/jeb.026880)

32. Eason EV, Hawkes EW, Windheim M, Christensen DL, Libby T, Cutkosky MR. 2015 Stress distribution and contact area measurements of a gecko toe using a high-resolution tactile sensor. Bioinspir. Biomim. 10, 016013. (doi:10.1088/1748-3190/10/1/016013)

33. Sharp JS, Poole SF, Kleiman BW. 2018 Optical measurement of contact forces using frustrated total internal reflection. Phys. Rev. Appl. 10, 034051. (doi:10.1103/physrevapplied.10.034051)

34. Weihmann T, Blickhan R. 2009 Comparing inclined locomotion in a ground-living and a climbing ant species: sagittal plane kinematics. J. Comp. Physiol. A 195, 1011–1020. (doi:10.1007/s00359-009-0475-y)

35. Reinhardt L, Blickhan R. 2014 Level locomotion in wood ants: evidence for grounded running. J. Exp. Biol. 217, 2358–2370. (doi:10.1242/jeb.098426)

36. Wöhrl T, Reinhardt L, Blickhan R. 2017 Propulsion in hexapod locomotion: how do desert ants traverse slopes? J. Exp. Biol. 220, 1618–1625. (doi:10.1242/jeb.137505)

37. Goldman DI, Chen TS, Dudek DM, Full RJ. 2006 Dynamics of rapid vertical climbing in cockroaches reveals a template. J. Exp. Biol. 209, 2990–3000. (doi:10.1242/jeb.02322)

38. Chun C, Biswas T, Bhandawat V. 2021 Drosophila uses a tripod gait across all walking speeds, and the geometry of the tripod is important for speed control. eLife 10, e65878. (doi:10.7554/eLife.65878)

