## Supplementary materials for "Quasi-static force requirements are not sufficient to explain arolium engagement in climbing Argentine ants"

### Supplementary material

**Method S1. Step Separation**

This section describes how steps were separated for each video.

All videos were first separated into strides and then divided into steps. Strides were defined as the time duration between one hind leg touchdown to its next touchdown. Over multiple strides, we identified the periodic lowest points of the foot-body distance and defined each minimum as the beginning of a stride. After tracking body and limb keypoints, each time series was segmented into strides.

We then separated the strides into stance phase and swing phase. The separation point is the first frame when the hind leg moves faster than 16mm/s. This empirical threshold was chosen because it is high enough to exceed noise levels yet low enough to represent the transition from stance to swing. From the beginning of the stride to the frame before the separation point was defined as the stance phase, and from the separation point to the end of the stride was defined as the swing phase.

We sought to only analyze climbing behaviors with a consistent alternating tripod gait, thus for every stride we calculated the tripod coordination strength (TCS). The TCS for both stance and swing phases was calculated using the same method described for step TCS in the main text. For each stride, the overall TCS was computed as the mean of the TCS values calculated separately for its stance and swing phases.

Only strides with its TCS greater than 0.5 were kept. As we were interested in the stance phases, we kept all stance phases for each stride. Because all retained strides had high tripod coordination strength (TCS > 0.5), indicating that the two tripod leg sets moved predominantly in alternation, we defined the swing phases as the stance phases for the alternate tripod leg set. To simplify the names, we called all the stance phases as steps.

**Method S2. Calculation of Arolium Engagement Intensity**

This section describes the procedure used to calculate arolium engagement intensity from video recordings.

1) **Background subtraction:** We first computed a background image by taking the median of a set of sampled frames. Frames were sampled every five frames across the entire video. The resulting background image was then subtracted from all frames to remove static visual features.
2) **Foot position extraction:** Tracking data obtained from DeepLabCut were used to determine the positions of all six feet in each frame. For each tracked foot position, we extracted a 25 × 25 pixel region centered on the foot location.
3) **Identify the maximum pixel value:** Within each 25 × 25 region, we found the point with maximum pixel value. If the maximum value exceeded the threshold T1 (the greater value between the median value of the 25 x 25 region plus 1400 and zero). A 5 × 5 pixel region centered on this point was then extracted. The 5 × 5 size was chosen because the arolium is smaller than this spatial window. Within the 5 × 5 region, we compared the maximum pixel value to the minimum pixel value plus 3000 (threshold T2). If the maximum value exceeded this threshold, the point with maximum pixel value is treated as an engaged point. The 1400 used in calculating threshold T1 and 3000 used in calculating threshold T2 were selected empirically.
4) **Check the 5 x 5 surrounding region:** We identified all pixels in the 5 × 5 region whose values were greater than threshold T1. Connected pixel clusters were then extracted using MATLAB's bwconncomp() function. The connected segment containing the maximum-value pixel was classified as the engaged region.
5) **Calculate the engagement intensity:** For each foot, the arolium engagement intensity was defined as the sum of the pixel values of all pixels within the identified engaged segment.

**Method S3. FTIR sensor validation**

This section describes the results of a supplementary experiment, which verified that the illuminated contacts originated specifically from the arolium and not from other leg structures.

In the supplementary experiment, we recorded the level walking of Argentine ants on our Frustrated Total Internal Reflection (FTIR) sensor, and compared the occurrence of the illuminated contact events with that observed in our climbing experiment. Illuminated contact events were rarely detected in level walking videos, occurring in only 0.5%–17.7% of strides across the 8 collections. In contrast, our climbing videos showed frequent illuminated contact, with 76.7%–91.6% of strides showing illumination across the 8 collections (Figure S3).

Previous research found that during level walking, weaver ants stood mainly on their 3rd and 4th tarsomeres, with tarsal hair instead of arolium touching the surface [1]. Conversely, ablation studies demonstrated that arolium engagement is essential for climbing smooth artificial surfaces [2]. Together, these findings verified that the FTIR system detected arolium engagement with high sensitivity but was largely unresponsive to tarsal hair contact, confirming that the illuminated contacts specifically represented arolium engagement.

**Method S4. 3D tripod quasi-static force model**

This section includes the details of the 3D tripod quasi-static force model.

As described in the main text, the quasi-static tripod model of climbing was constructed as a point mass with three weightless limbs representing the ant's center of mass connected with the 3 stance phase limbs. The center of mass (CoM) is located a distance $h$ from the surface in the z direction, and the limb locations in the xy-plane are measured from CoM. The mass point is subjected to gravity ($m\boldsymbol{g}$), while 3-dimensional foot contacting forces ($\boldsymbol{F}_1, \boldsymbol{F}_2, \boldsymbol{F}_3$) act on the tip of all stance-phase limbs (Figure S4). To maintain the quasi-static force balance, the sum of gravitational ($m\boldsymbol{g}$) and foot contact forces ($\boldsymbol{F}_1, \boldsymbol{F}_2, \boldsymbol{F}_3$) must be zero in all three directions, and the sum of torques about the center of mass due to the foot contact forces ($\boldsymbol{\tau}_1,\ \boldsymbol{\tau}_2,\ \boldsymbol{\tau}_3$) must be zero for all three axes:

$$\sum_{i=1}^{3} \boldsymbol{F}_i + m\boldsymbol{g} = 0 \Rightarrow \begin{cases} F_{1x} + F_{2x} + F_{3x} = mg \\ F_{1y} + F_{2y} + F_{3y} = 0 \\ F_{1z} + F_{2z} + F_{3z} = 0 \end{cases} \tag{1}$$

$$\sum_{i=1}^{3} \boldsymbol{\tau}_i = 0 \Rightarrow \begin{cases} y_1F_{1Z} + y_2F_{2Z} + y_3F_{3Z} - h(F_{1y} + F_{2y} + F_{3y}) = 0 \\ -(x_1F_{1Z} + x_2F_{2Z} + x_3F_{3Z}) - h(F_{1x} + F_{2x} + F_{3x}) = 0 \\ x_1F_{1y} + x_2F_{2y} + x_3F_{3y} - (y_1F_{1x} + y_2F_{2x} + y_3F_{3x}) = 0 \end{cases} \tag{2}$$

Solving equations (1) and (2), yields the expression for $F_{1z},\ F_{2z},\ F_{3z}$ in the main text.

$$F_{1z} = \frac{mgh(y_3 - y_2)}{D} \tag{3}$$

$$F_{2z} = \frac{mgh(y_1 - y_3)}{D} \tag{4}$$

$$F_{3z} = \frac{mgh(y_2 - y_1)}{D} \tag{5}$$

$$D = (x_1 - x_3)(y_2 - y_3) - (x_2 - x_3)(y_1 - y_3) \tag{6}$$

Where $F_{1x},\ F_{2x},\ F_{3x}$ represent the signed scalar fore-aft components, $F_{1y},\ F_{2y},\ F_{3y}$ represent the signed scalar lateral components and $F_{1z},\ F_{2z},\ F_{3z}$ represent the signed scalar normal components of the foot contact forces. $mg$ represents the magnitude of the gravitational force acting on the object, $h$ is a positive scalar representing the distance between the point mass and the contacting surface. $x_1,\ x_2,\ x_3$ and $y_1,\ y_2,\ y_3$ correspond to the x and y signed scalar components of the foot contact points relative to the coordinate origin. The positive directions for force components ($F_{ix}$,

$F_{iy}$, $F_{iz}$) and position coordinates ($x_i$, $y_i$) are indicated in Figure S4 by colored arrows: red for fore-aft, green for lateral and blue for normal. The foot numbering is arbitrary and does not influence the results.

Ant anatomy restricts the possible directions of the quasi-static normal forces in the tripod model to just two distinct patterns, as shown in Figure 2 in the main text. Without loss of generality, we numbered feet 1–3 sequentially from uppermost to lowest along the gravity vector. During upward climbing, the uppermost foot is the front foot and the lowest foot is the hind foot. During downward climbing, this relationship reverses: the uppermost foot is the hind foot and the lowest foot is the front foot.

Anatomically, the fore-aft distance between the uppermost and the lowest feet exceeds that between the middle and the lowest feet. During a tripod stance, the uppermost and the lowest feet lie on the same lateral side of the center of mass, while the middle foot is contralateral. Consequently, lateral distances are smaller between the uppermost and the lowest feet than that between the middle and the lowest feet. This geometry results in the inequality $|(x_1 - x_3)(y_2 - y_3)| > |(x_2 - x_3)(y_1 - y_3)|$. Consequently, the sign of the denominator $D$ (equation 6) is determined by the first cross product $(x_1 - x_3)(y_2 - y_3)$. Since the fore-aft term $(x_1 - x_3)$ is positive, the sign of $D$ depends solely on the lateral offset $(y_2 - y_3)$.

As $F_{1z}$'s numerator $(y_3 - y_2)$ has the opposite sign and $F_{3z}$'s numerator $(y_2 - y_1)$ has the same sign of this lateral offset, the model predicts $F_{1z}$ is negative (adhesion force) and $F_{3z}$ is positive (compression force). The normal force on the middle foot can be either adhesive or compressive, depending on the sign of the lateral offset between the uppermost and the lowest feet $(y_1 - y_3)$. This sign corresponds to two geometric configurations, which we classify as distinct stance types (wide stance and narrow stance) in the main text.

**Figure S1:**

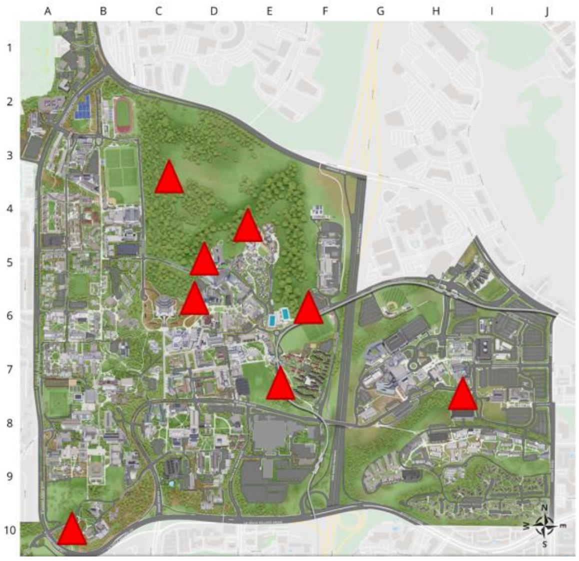


**Figure S1: Ant collecting locations.** The background is a UC San Diego campus map and the red triangles represent the eight locations where ants in this experiment were collected.

**Figure S2:**

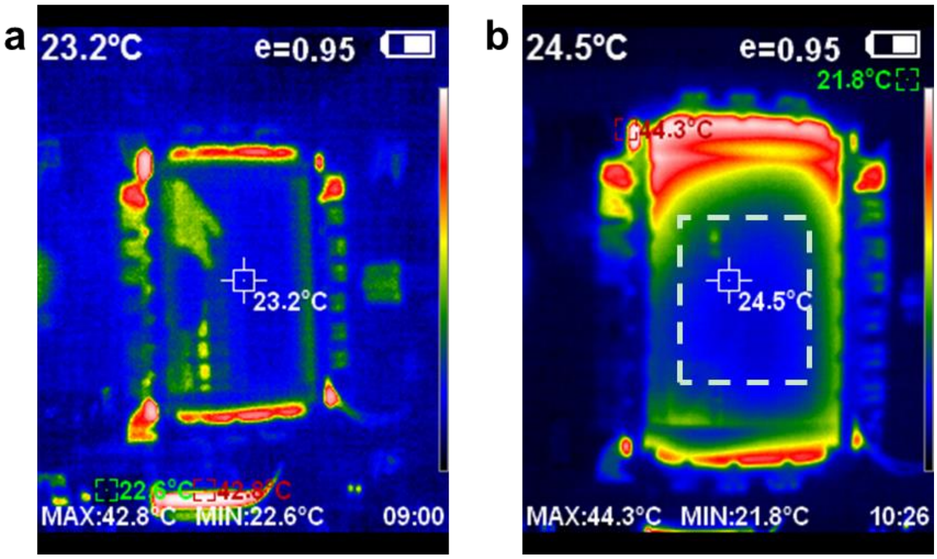


**Figure S2: Thermograms of the FTIR plate. (a)** Thermogram taken when the Led strip and the fan just turned on. **(b)** Thermogram taken 1 hour and 26 minutes after **(a)**. The region inside the dashed white line was the area accessible to ants. It took less than an hour for the FTIR temperature to reach a steady state. During the steady state, the ant walking area is in the temperature range of 22~34°C with most of the area staying around 24°C

**Figure S3:**

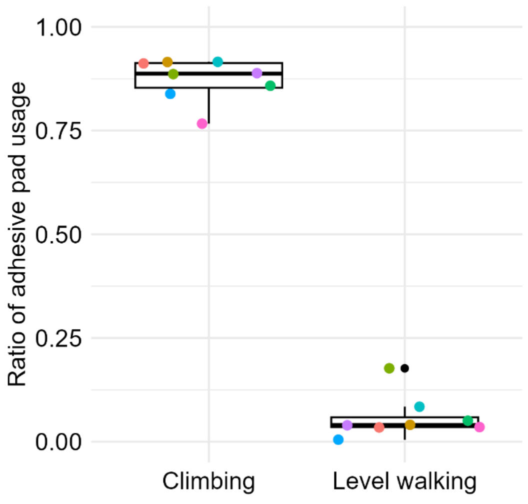


**Figure S3: Ratio of adhesive pad usage.** The ratio is the number of strides with illuminated contact divided by the total number of strides. Dots in different colors represent the ratios of different collections. The number of strides for climbing varied from 238 to 4193 across collections, and the number of strides for level walking varied from 462 to 3576 strides across collections.

**Figure S4:**

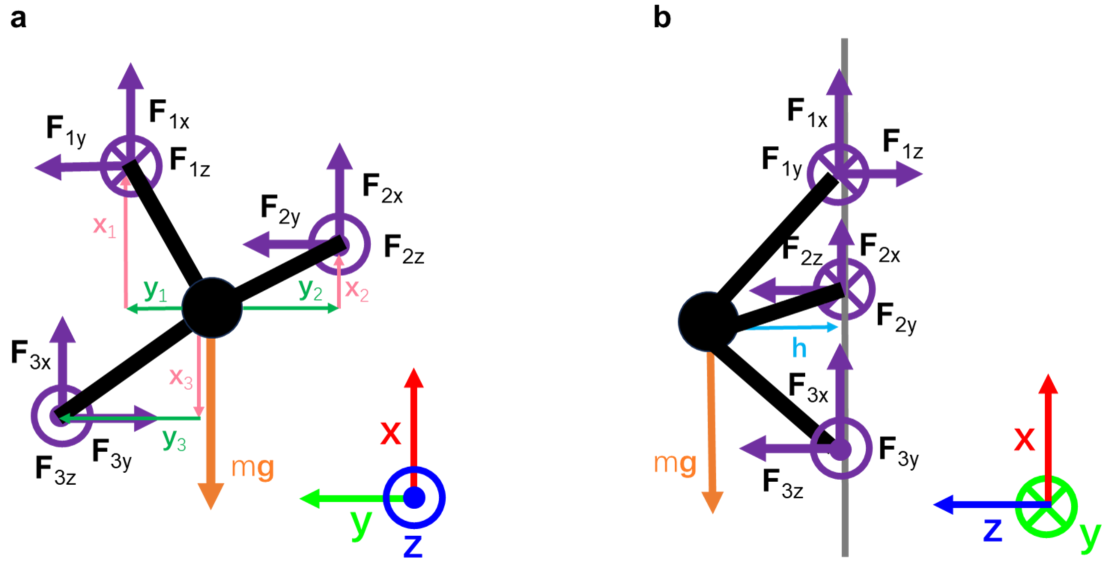


**Figure S4: Tripod model diagram.** (**a**) Dorsal view of the tripod model. (**b**) Lateral view of the tripod model. The ant was represented as a black mass point with three black, massless limbs. The gray vertical line in (**b**) represents the glass surface. Foot contact forces were depicted by purple arrows ($\boldsymbol{F}_{1x,1y,1z}$, $\boldsymbol{F}_{2x,2y,2z}$, $\boldsymbol{F}_{3x,3y,3z}$) and the gravitational force was shown as an orange arrow ($m\boldsymbol{g}$). Red, green, and blue arrows in the bottom right corner represent the fore-aft ($\boldsymbol{x}$), lateral ($\boldsymbol{y}$), and normal directions ($\boldsymbol{z}$), respectively, with arrows pointing in the positive direction of each axis. $\otimes$ denotes an arrow directed into the page, while $\odot$ denotes an arrow directed out of the page. The center of mass (CoM) is located a distance $h$ from the surface in the z direction, labeled by a light blue arrow ($\boldsymbol{h}$) in (**b**), and the foot locations in the xy-plane are measured from CoM, with their $\boldsymbol{x}$ ($\boldsymbol{x}_1, \boldsymbol{x}_2, \boldsymbol{x}_3$) and $\boldsymbol{y}$ ($\boldsymbol{y}_1, \boldsymbol{y}_2, \boldsymbol{y}_3$) vector components represented by pink and darker green arrows in (**a**), respectively.

#### Figure S5:

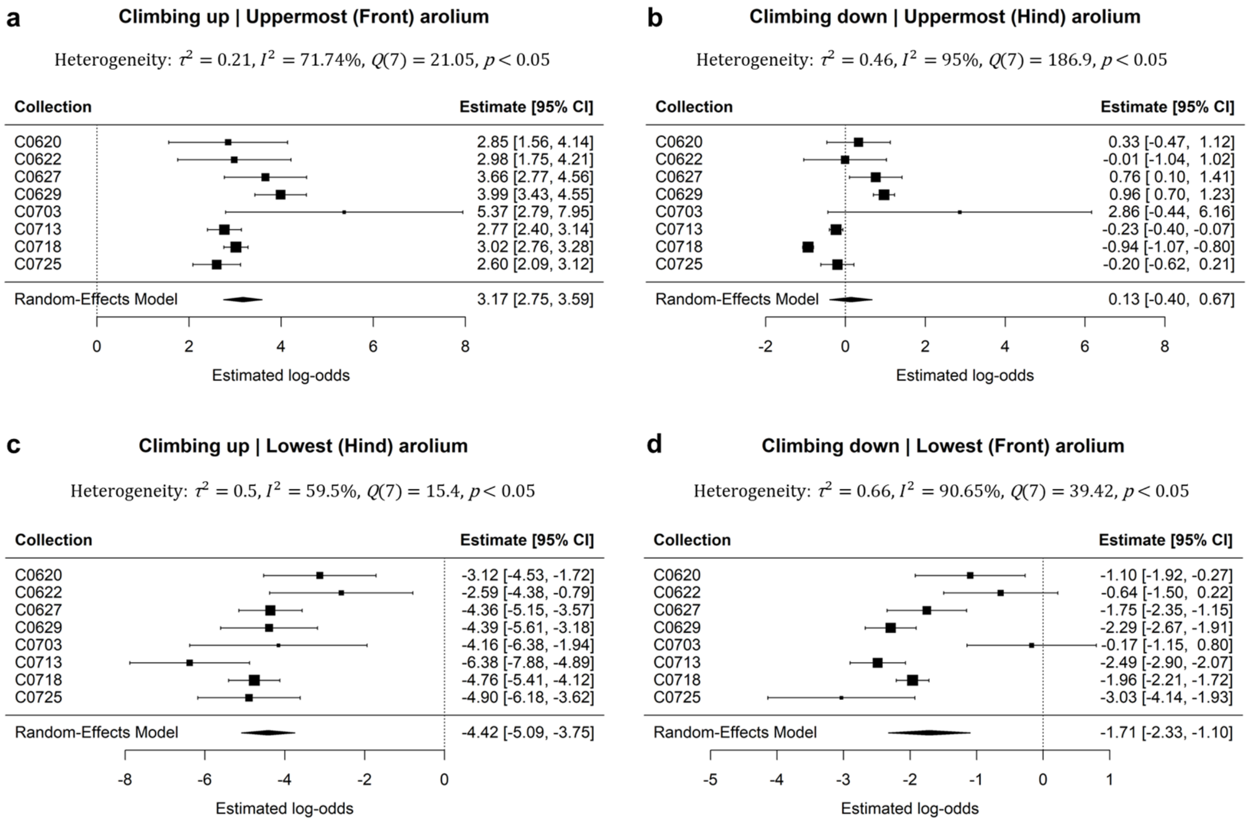


**Figure S5: Forest plots showing estimated log-odds of engagement for the uppermost and lowest arolia.** Uppermost arolium during upward climbing (**a**) and downward climbing (**b**). Lowest arolium during upward climbing (**c**) and downward climbing (**d**). Each point represents the estimated log-odds of engagement from a single collection, and the corresponding error bar indicates the corresponding 95% confidence interval. The black diamond shows the pooled log-odds estimate and its 95% confidence interval of all collections.

**Figure S6:**

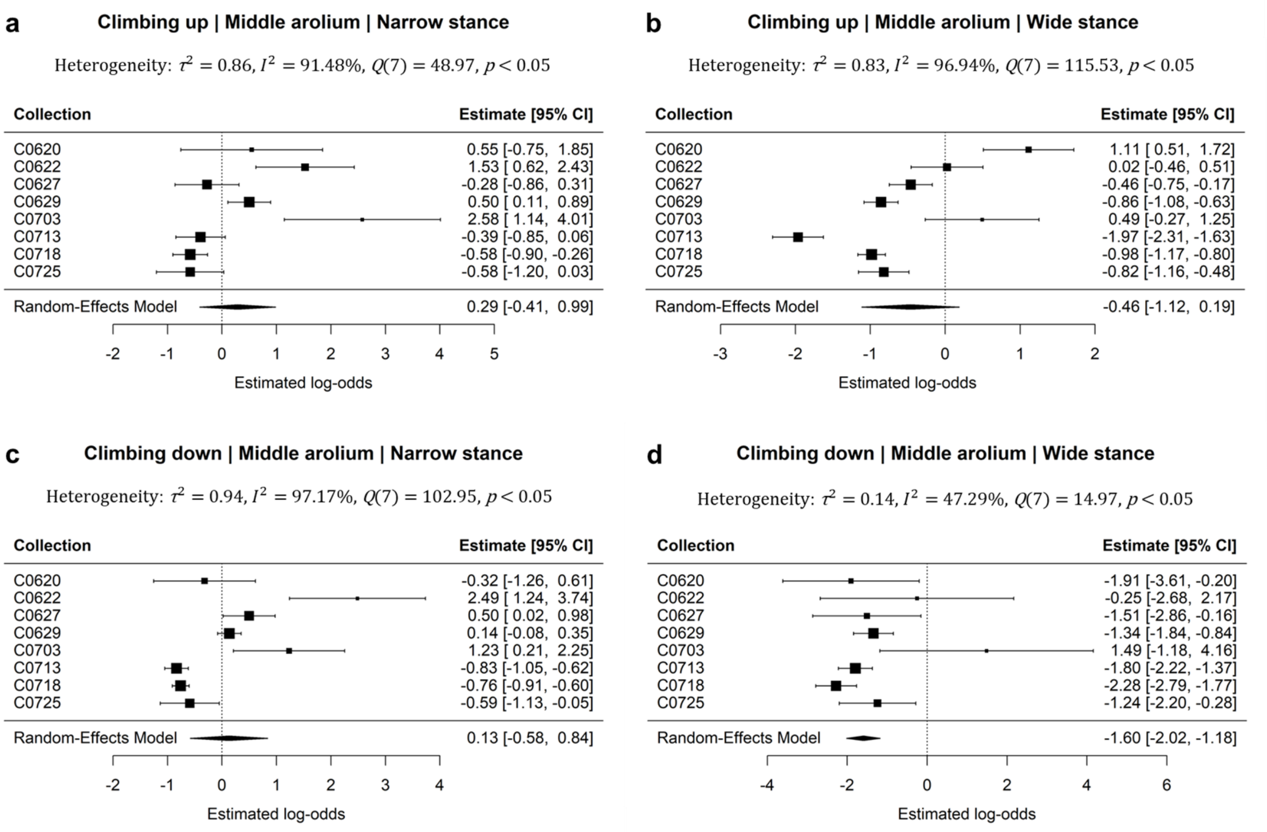


**Figure S6: Forest plots showing estimated log-odds of engagement for the middle arolium.** Middle arolium during upward climbing in narrow (**a**) and wide (**b**) stances. Middle arolium during downward climbing in narrow (**c**) and wide (**d**) stances. Each point represents the estimated log-odds of engagement from a single collection, and the corresponding error bar indicates the corresponding 95% confidence interval. The black diamond shows the pooled log-odds estimate and its 95% confidence interval of all collections.

**Figure S7:**

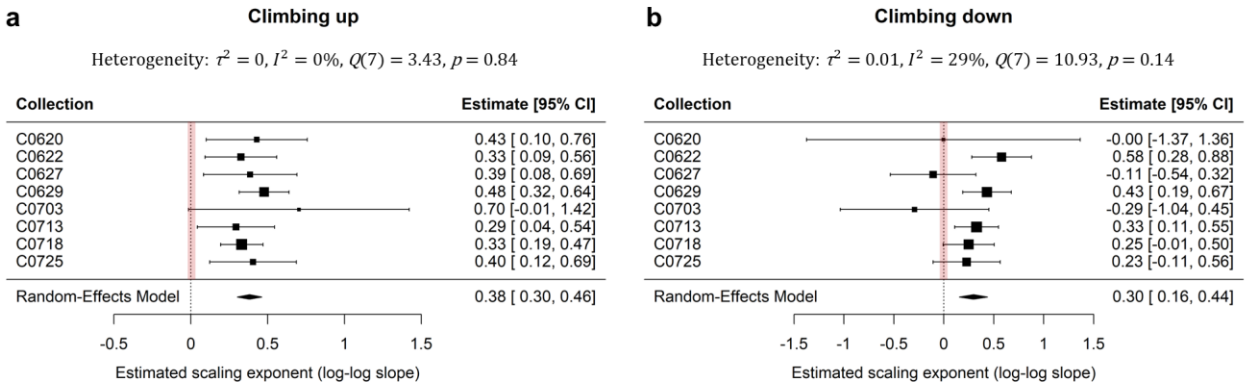


**Figure S7: Forest plots showing estimated scaling exponents between the engagement intensity ratio and the required adhesion force ratio.** Estimated scaling exponents during upward (**a**) and downward (**b**) climbing. Each point represents the estimated scaling exponent from a single collection, and the corresponding error bar indicates the corresponding 95% confidence interval. The black diamond shows the pooled scaling exponent estimate and its 95% confidence interval of all collections. The vertical dashed line with red shade at scaling exponent = 0 indicates no relationship between engagement intensity ratio and required adhesion force ratio. Negative exponents indicate engagement intensity ratio decreasing with the required adhesion force ratio, whereas positive exponents indicate intensity ratio increasing with the force ratio.

**Figure S8:**

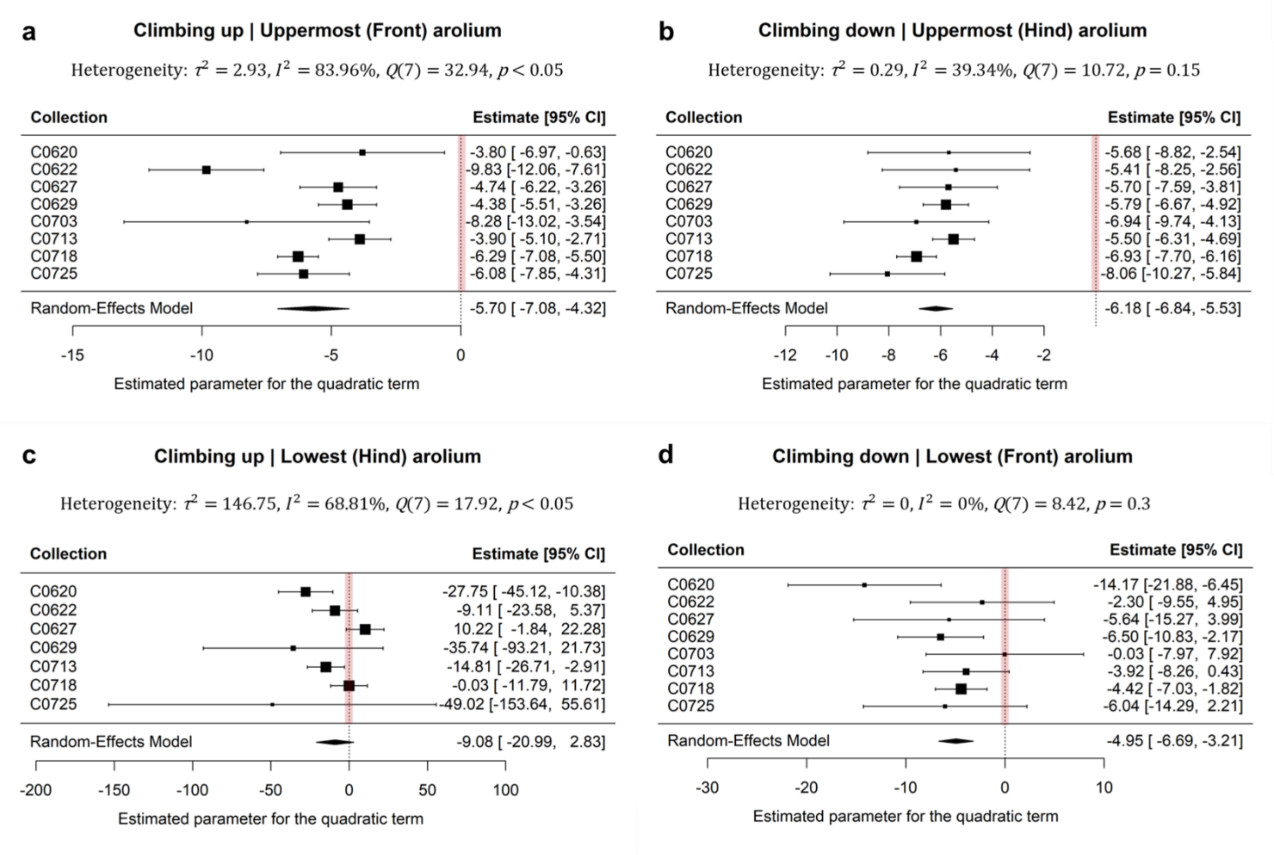


**Figure S8: Forest plots of estimated quadratic coefficients for the uppermost and lowest arolia.** Uppermost arolium during upward climbing (**a**) and downward climbing (**b**). Lowest arolium during upward climbing (**c**) and downward climbing (**d**). The model is $y = a(x - 0.5)^2 + b(x - 0.5) + c$, where $y$ is the standardized arolium engagement intensity, $x$ is the normalized time, $a$ is the quadratic coefficient, $b$ is the linear coefficient, and $c$ is the intercept. Each point represents the estimated parameter for the quadratic term from a single collection, and the corresponding error bar indicates the corresponding 95% confidence interval. The black diamond shows the pooled parameter estimate for the quadratic terms and its 95% confidence interval of all collections. Data to the left of the vertical dashed line with red shade (negative values) indicate a peak-shaped pattern.

**Figure S9:**

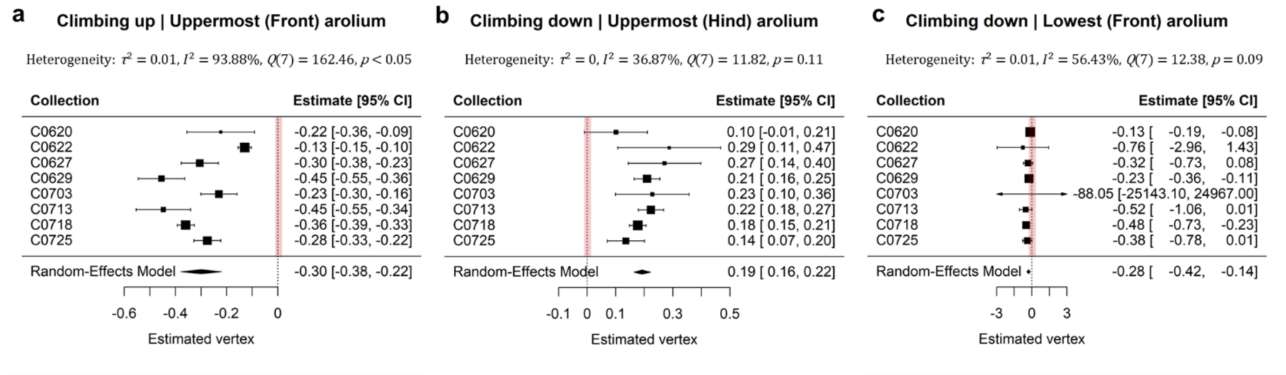


**Figure S9: Forest plots of estimated vertex.** Uppermost arolium during upward (**a**) and downward climbing (**b**). Lowest arolium during downward climbing (**c**). Each point represents the estimated vertex ($-\frac{b}{2a}$) from a single collection, and the corresponding error bar indicates the corresponding 95% confidence interval. The black diamond shows the pooled vertex estimate and its 95% confidence interval of all collections. The vertical dashed line with red shade at vertex = 0 indicates a symmetric pattern peaking at mid-stance. Negative values indicate an asymmetric pattern peaking before mid-stance, whereas positive values indicate an asymmetric pattern peaking after mid-stance. Vertices reported in the main text are these values plus 0.5.

**Figure S10:**

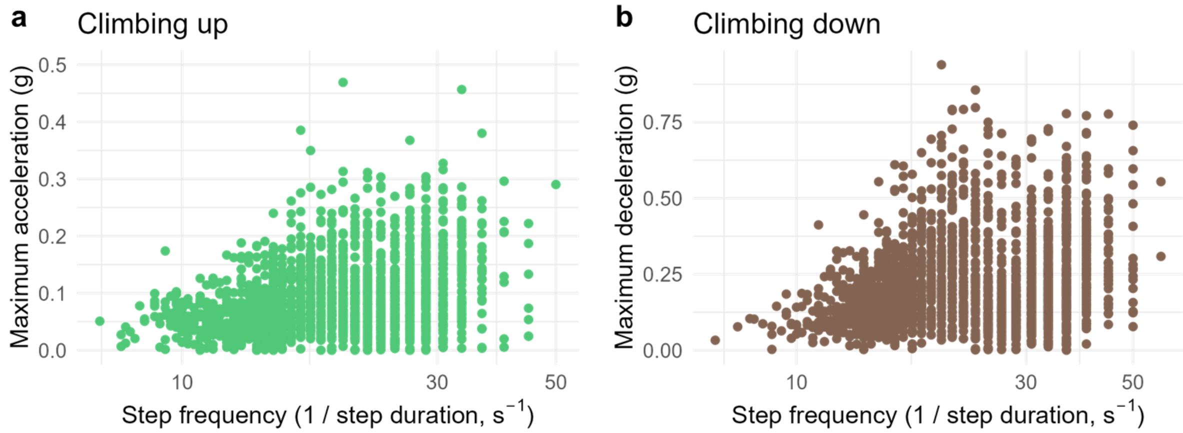


**Figure S10: Maximum acceleration/deceleration during each step versus step frequency.** (**a**) Maximum acceleration versus step frequency during upward climbing. (**b**) Maximum deceleration versus step frequency during downward climbing. During upward climbing, we analyzed the acceleration phase, whereas during downward climbing, we analyzed the deceleration phase, as both require higher-than-normal leg forces in the normal direction to prevent detachment. g is $9.8 m/s^2$.

**Figure S11:**

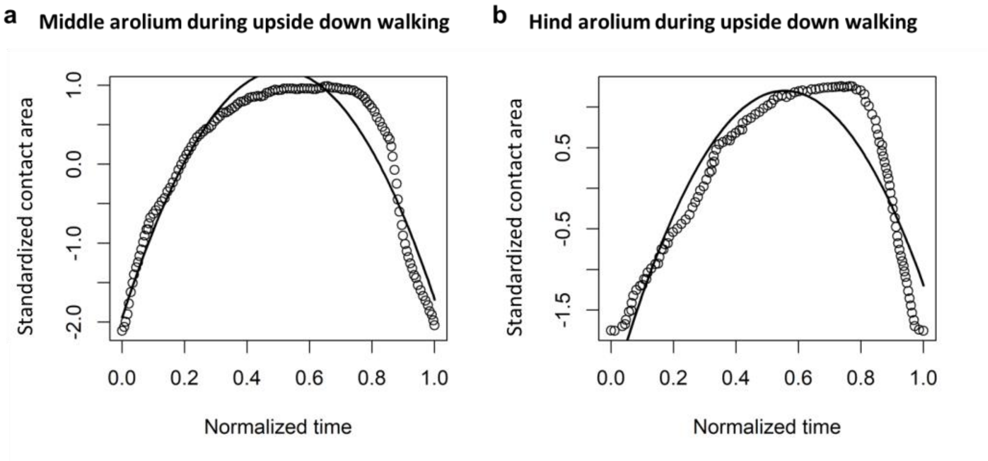


**Figure S11: Temporal variation of contact area during upside-down walking.** (**a**) Middle arolium. The circular points represent a standardized version of the data collected from Figure 2*c* in [3]. (**b**) Hind arolium. The circular points represent a standardized version of the data collected from Figure 7 in [3]. Curves in (**a**) and (**b**) are fitted polynomial regressions using the same model as in Figure S8.

### References


1. Endlein T, Federle W. 2015 On heels and toes: how ants climb with adhesive pads and tarsal friction hair arrays. *PLoS ONE* **10**, e0141269. (doi:10.1371/journal.pone.0141269)
2. Brechka P. 2024 Slippery surfaces and the biomechanics of climbing in Macaranga-Ant mutualisms. PhD thesis. Cambridge, UK: University of Cambridge.
3. Federle W, Endlein T. 2004 Locomotion and adhesion: dynamic control of adhesive surface contact in ants. *Arthropod Struct. Dev.* **33**, 67–75.
